# Scube2 Modulates Coronary Vessel Formation during Cardiac Growth and Regeneration in Zebrafish

**DOI:** 10.64898/2025.12.17.695032

**Authors:** Ann Nee Lee, Ke-Hsuan Wei, Kaushik Chowdhury, Muhammad Abdul Rouf, An-Ju Chu, Yu-Jen Hung, Ku-Chi Tsao, Yan-Ting Chen, Yuh-Charn Lin, Yao-Ming Chang, Rubén Marín-Juez, Ruey-Bing Yang, Shih-Lei (Ben) Lai

## Abstract

Coronary vessel formation is essential for cardiac growth and regeneration, yet the mechanisms regulating these processes remain incompletely defined. In this study, we identified and characterized Scube2 as a novel regulator in coronary vessel formation and heart regeneration. Scube2 is expressed in the epicardium covering the coronary vasculature under homeostatic conditions in adult hearts and is strongly upregulated after cardiac injury. Zebrafish *scube2* mutant exhibits reduced coronary vessel coverage during cardiac growth and pronounced myocardial defects, including abnormal vessel morphology, disorganized contractile fibers, and mitochondrial damage. Following injury, *scube2* mutant hearts show impaired revascularization and persistent fibrosis, indicating defective regeneration. Mechanistically, transcriptomic profiling of *scube2* mutant hearts reveals attenuated expression of angiogenic-related genes, including those mediated by vascular endothelial and platelet-derived growth factors, consistent with reduced proliferation of endothelial cells and cardiomyocytes during repair. In a gain-of-function setting, SCUBE2 enhances platelet-derived growth factor signaling by promoting receptor activation in HEK293 cells. These findings support a novel role for Scube2 in cardiac growth and regeneration by modulating coronary vessel formation.

**Graphical Abstract:** 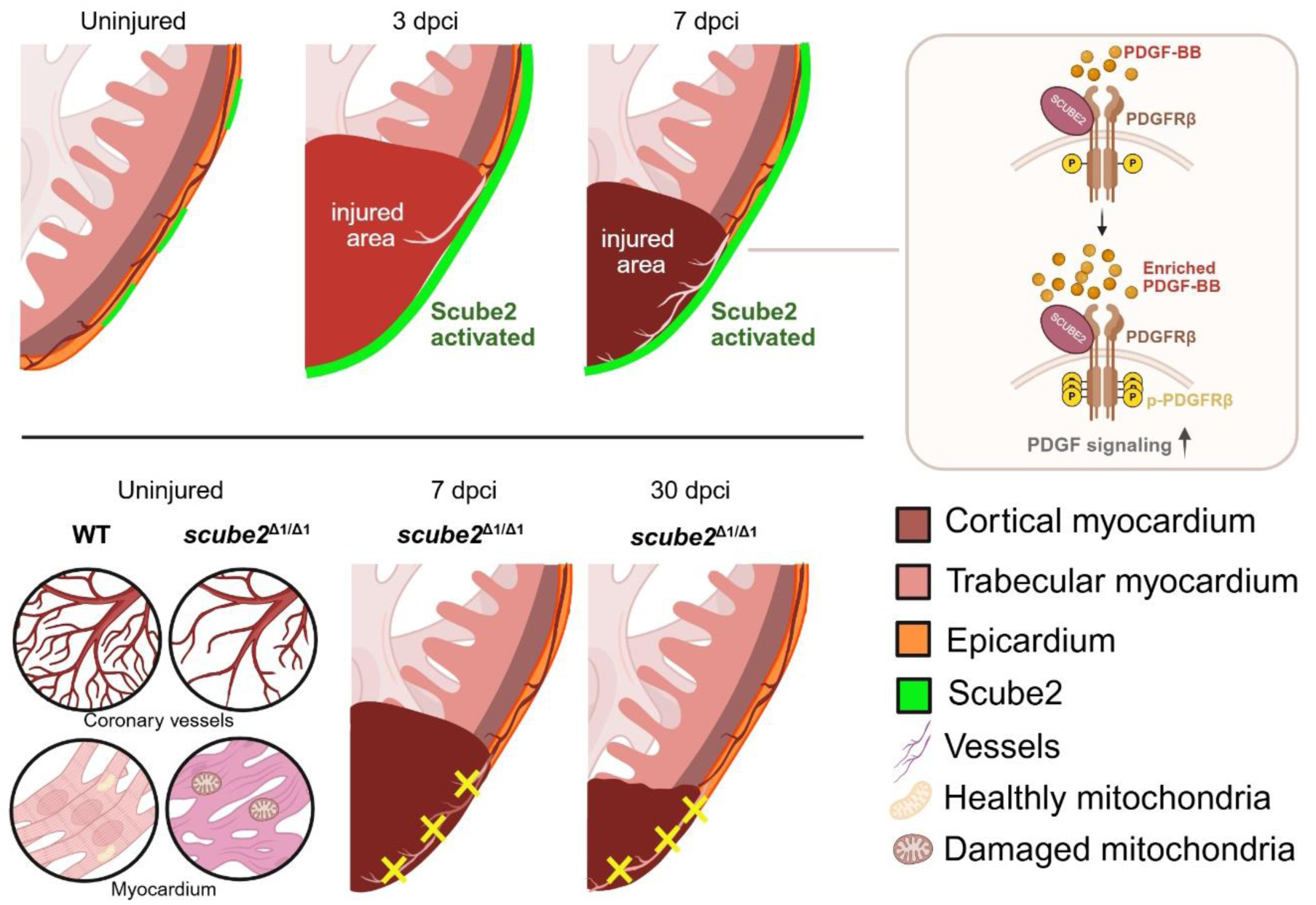

## Introduction

The coronary vasculature is essential for supplying oxygen and nutrients to the thickening myocardium during heart growth and for revascularization during cardiac repair after myocardial infarction (MI). In mammals, coronary vessels develop through angiogenic sprouting and remodeling, ultimately forming a hierarchical network of arteries, veins, and capillaries (1–4). This vascular network continues to expand postnatally to support myocardial maturation (1). In parallel, pathological occlusion of coronary arteries, such as in coronary artery disease (CAD), leads to MI, resulting in cardiomyocyte (CM) death and heart failure (5, 6). While therapeutic revascularization strategies aim to restore perfusion, clinical translation remains limited by an incomplete understanding of the underlying regulatory mechanisms (7, 8).

Vascular endothelial growth factor (VEGF) signaling is indispensable for coronary vessel formation and post-injury angiogenesis (9, 10). Among potential regulators of VEGF signaling, recent studies have identified the signal peptide-CUB (complement C1r/C1s, Uegf, Bmp1)-EGF (epithelial growth factor) domain-containing protein 2 (SCUBE2), a secretory glycoprotein, as a co-receptor of VEGFR2 when tethered to the plasma membrane of endothelial cells and regulates their proliferation and tube formation (11).

The zebrafish Scube2 homolog also enhances VEGF signaling during embryonic development, promoting endothelial proliferation, migration, and tube formation (12). Besides VEGF, Scube2 may also function non-cell autonomously in regulating other signaling pathways, including long-range hedgehog signaling during zebrafish embryonic development (13–15). However, its potential role in coronary development and regeneration in the adult heart remains largely unknown.

The zebrafish is a powerful model to dissect the molecular mechanisms of cardiovascular development and regeneration (16). Zebrafish hearts exhibit conserved structural, genetic, and physiological features similar to those of humans, including the development of coronary vessels (17, 18). In contrast to mammals, zebrafish can fully regenerate their hearts after injury through rapid revascularization, cardiomyocyte proliferation, and scar resolution (9, 19–21). Notably, revascularization of the injured area is one of the first events induced by cardiac injury and is essential for heart regeneration (9, 18, 22).

Here, we investigate the role of Scube2 in coronary vessel formation and cardiac regeneration using zebrafish models. We employ genetic knockout, high-resolution imaging, and transcriptomic profiling to examine the impact of Scube2 on coronary growth, post-injury revascularization, and heart regeneration. Our study reveals a previously unrecognized function of Scube2 in regulating coronary vascularization and CM proliferation, highlighting its potential as a therapeutic target for ischemic heart disease.

## Methods

### Ethics statement

Zebrafish were handled according to the Institutional Animal Care and Utilization Committee of Academia Sinica (Protocol 21-12-1749), anesthetized with tricaine methanesulfonate (MS-222, 0.02%), and euthanized with MS-222 (0.05%) following established protocols. All euthanasia procedures were consistently applied to the respective experimental groups.

### Zebrafish lines and maintenance

Wild-type AB strain, *scube2^Δ1/Δ1^* mutant (*as403*), and transgenic zebrafish *Tg(fli1:EGFP)^y1^*(23), *Tg(myl7:nuc-RFP)^f2^*(24)*, Tg(tcf21:NLS-EGFP)^pd41^*(25)*, Tg(wt1b:EGFP)^li1^* (26), were maintained in a 14-h light/10-h dark cycle at 28.5°C following standard protocol. *Scube2^Δ1/Δ1^* mutants were generated as described (27), and mutants used in this study were produced by homozygous mutant incross.

### Antibodies and reagents

Anti-FLAG (catalog F7425) antibody was from Sigma-Aldrich (St. Louis, MO, USA). Anti-HA (catalog 3724), anti-phospho-PDGFRβ (Tyr751) (catalog 3161), and anti-PDGFRβ (catalog 3169) antibodies were from Cell Signaling Technology (Boston, MA, USA). Horseradish peroxidase (HRP)-conjugated secondary antibody was from Jackson ImmunoResearch Laboratories (West Grove, PA, USA). Anti-His antibody used in this study was designed and produced in-house by our laboratory. Recombinant human PDGF-BB protein (rhPDGF-BB) (catalog 220-BB) was from R&D Systems (Minneapolis, MN, USA). Lipofectamine 3000 was from Thermo Fisher Scientific (Waltham, MA, USA).

### Coronary Vessel Imaging and Quantification

Coronary vasculature was examined using *Tg(fli1:EGFP)^y1^* transgenic background in combination with confocal microscopy. Fish at defined developmental stages (17 mm and 28 mm snout-fin base length) were euthanized and heart extracted. The ventricles were then placed into 1X PBS and rinsed several times for at least 3 minutes, followed by 4% paraformaldehyde (PFA) treatment for at least 5 minutes. The ventricles were mounted in a bottom glass slide with 1% low melting agarose before imaging with Leica SP8/SP8 DLS Confocal microscopy with 10X magnification and Z-stack 3 (17 mm snout-fin base length) and 4 (28 mm snout-fin base length). Quantification of vessel coverage was performed using ImageJ by calculating the ratio of EGFP-positive vessel area to ventricular surface area.

### Cardiac Injury

Cryoinjury was performed as previously described in adult zebrafish (6-12 months old)(20, 21). In brief, fish were anesthetized in 0.04% tricaine (Sigma, St Louis, MO) and immobilized in a damp sponge with the ventral side up. A small incision was created through the thoracic wall by microdissection scissors. A stainless steel cryoprobe precooled in liquid nitrogen was placed on the ventricular surface until it thawed. Fish were then returned to a freshwater tank for recovery, and their reanimation could be enhanced by gently pipetting water onto the gills after surgery.

### Cryosections and histological analyses

Zebrafish hearts were extracted and fixed in 4% (wt/vol) paraformaldehyde (Alfa Aesar, MA) at room temperature for 1 hr. Collected hearts were subsequently cryopreserved with 30% (wt/vol) sucrose, followed by immersion in OCT (Tissue-Tek, Sakura Finetek, Torrance, CA) and stored at −80°C immediately. 10 μm cryosections were collected for histological analysis. AFOG staining was applied for the visualization of healthy CMs in orange, collagens in blue, and fibrins in red. In brief, samples were incubated in preheated Bouin’s solution (Sigma) at 58°C for 2 hours post-fixation and subsequently immersed in a 1% phosphomolybdic acid (Sigma) and 0.5% phosphotungstic acid solution (Sigma) at room temperature for 5 minutes for mordanting. Samples were then incubated with AFOG solution containing Aniline Blue (Sigma), Orange G (Sigma), and Acid Fuchsin (Sigma) for color development. Quantification was done by measuring the scar area in each heart from five discontinuous sections, including the one with the largest scar, as well as the two sections at the front and the two sections at the back, as previously described (28). Statistical analysis was performed using Prism 9, with one-way ANOVA.

### Immunostaining and Cell Proliferation Assay

For immunofluorescence, slides were washed twice with PBS and twice with ddH2O, underwent antigen retrieval with 10 mM citrate buffer, then incubated in blocking solution (1×PBS, 2% [vol/vol] sheep serum, 0.2% Triton X-100, 1% DMSO). Slides were then incubated in primary antibodies overnight at 4°C, followed by three washes in PBST (0.1% Triton X-100 in 1× PBS) and incubation with secondary antibodies for 1.5 hours at 28°C. Slides were washed again with PBST and stained with DAPI (Santa Cruz Biotechnology, TX) before mounting. Antibodies and reagents used in this study included anti-mCherry (Abcam, Cambridge, UK) at 1:250, anti-fli1 (Abcam, Cambridge, UK) at 1:200, and anti-PCNA (Santa Cruz Biotechnology, TX) at 1:250. Imaging of whole-mount hearts and heart sections was performed using Nikon SMZ25 and Zeiss LSM 780, respectively. Proliferating CM and the density of CM nuclei in the 200 μm border zone directly adjacent to the injured area were quantified as previously described(9). Proliferating cEC and the density of cEC nuclei in the injury zone were quantified as previously described (Fli1+/PCNA+/DAPI+) (18). Revascularization was examined by live imaging of the endogenous fluorescence from vessel reporter fish. Revascularized vessel density in the whole injured area was measured and quantified using ImageJ. One-way ANOVA was applied to assess all comparisons using Prism 9.

### *In situ* hybridization

*In situ* hybridization was performed on cryosections according to standard procedures. Briefly, the template for the antisense digoxigenin (DIG)-labeled riboprobe for scube2 was generated by PCR with the additional T7 promoter sequence included in the primers. Oligonucleotide sequences for probe synthesis are as follows: *scube2_201_SP6F* AGACTGTATCGAGGCAGA and *scube2_201_T7R* TCGGACGAGTTTCTTAGC. The PCR product was transcribed using DIG RNA labeling mix (Roche, Basel, Switzerland) and T7 RNA polymerase (Promega, WI). To prepare the cryosections, hearts were fixed in 4% PFA in diethyl pyrocarbonate (DEPC)-treated PBS at 4°C overnight and then immersed in OCT as described in the section ‘Cryosections and histological analyses.’ Briefly, 18 μm cryosections were used and dried at 37°C for 30 min, treated with 10 μg/ml proteinase K (Sigma) and post-fixed with 4% PFA for 15 min. Cryosections were prehybridized for at least 2 hr without probe and then hybridized with probes at 65°C overnight. After serial washing with 1×SSC and 1×MABT, the cryosections were blocked for 1 hr at room temperature and then incubated with anti-Digoxigenin Fab fragments Antibody, AP Conjugated (1:1000; Roche, Basel, Switzerland) at 4°C overnight. The next day, the probes were detected by chromogenic reaction with NBT/BCIP stock solution (1:50; Roche) and observed by a slide scanner (Pannoramic 250 FALSH II).

### Transmissible Electron Microscopy

The TEM procedures followed the protocols of a previous publication (29). Briefly, the ventricles were extracted and kept overnight at 4°C in fixative (2.5% glutaraldehyde in 1×PBS; pH: 7.4) and thereafter post-fixed in 1% OsO4 in the same buffer. After dehydration, in graded ethanol, the samples were finally embedded in Spurr resin (Spurr Low Viscosity Embedding Kit; EMS ®). Toluidine blue stain was used for staining semi-thin sections (500 nm) for light microscopy to guide EM selection. Subsequently, ultrathin sections were cut on a Leica® Ultracut UC7 Ultramicrotome (Leica Microsystems, Vienna, Austria) equipped with a diamond knife, stained with uranyl acetate and lead citrate then examined in a FEI Tecnai G2 F20 S-TWIN TEM (FEI, Hillsboro, OR, USA) at 120 kV in the Institute of Cellular and Organismic Biology, Academia Sinica.

### Transcriptomic Analysis

RNA was extracted from the hearts of uninjured and injured zebrafish at 3 and 7 days post-cryoinjury (dpci) in wild-type and scube2 mutant backgrounds. For each time point and condition, zebrafish ventricles from three fish hearts were used as biological duplicates for the RNA-seq experiments. RNA extraction was performed as previously described with minor modifications (28). Briefly, RNA isolation was done using the miRNeasy micro Kit (QIAGEN). RNA quality analysis was performed using Qubit and Bioanalyzer at the NGS High-Throughput Genomics Core, Academia Sinica. Sequencing was performed on the HiSeq Rapid (Illumina) setup, yielding 15–20 million reads per library on a 2 × 150 bp paired-end setup at the NGS core facility. Raw reads were assessed after quality control (QC) and output to filtered reads that were mapped to the zebrafish Ensembl genome assembly using HISAT2 (30). The number of reads was aligned to the Ensembl annotation using StringTie (31) to calculate gene expression as raw read counts. The raw counts of the mapped annotated genes were combined into a single matrix and normalized using upper-quantile normalization to generate normalized FPKM values. Further analysis was conducted based on the normalized read values and respective log2 fold changes to identify differentially expressed genes (DEGs) using NOIseq (32). GO enrichment analysis was performed in WebGestalt (33) for the dataset considering the upregulated and downregulated genes against the wild-type versus *scube2* mutant background. Over-representation analysis was done with an id of mapped genes for multiple testing corrections using Benjamini and Hochberg FDR correction, and conducted with a significance level of 0.05. PCA was performed on the normalized FPKM values of all the datasets at respective time points to analyze the sample level progression through time under control versus treatment conditions. Pathway analysis was performed using IPA software (QIAGEN)(34) following the manufacturer’s protocols. Log2 ratio was input from the normalized read counts in zebrafish and defined as DEGs at log2FC above ±1.5.

### Statistical Analysis

Data are presented as mean ± SEM. Statistical significance was assessed using Student’s t-test or ANOVA with appropriate post-hoc tests, as indicated. A p-value < 0.05 was considered statistically significant.

### Cell culture and transfection

Human embryonic kidney (HEK)-293T cells were obtained from the American Type Culture Collection (ATCC, Manassas, VA, USA; Catalog No. CRL-3216). Cells were authenticated by the supplier and routinely tested negative for mycoplasma contamination. All experiments involving human-derived cell lines were conducted in accordance with institutional biosafety regulations. HEK-293T cells were maintained in Dulbecco’s Modified Eagle Medium (DMEM) supplemented with 10% heat-inactivated fetal bovine serum, 100 units/ml penicillin, and 100 µg/ml streptomycin at 37°C in a 5% CO₂ atmosphere. The FLAG epitope-tagged SCUBE2 expression plasmid was prepared as previously described¹. The His-tagged PDGF-B (catalog HG10572-NH) and HA-tagged PDGFRβ (catalog HG10514-NY) expression plasmids were obtained from Sino Biological (Beijing, China). Cells were transfected with the plasmids using Lipofectamine 3000 for 48 hours. For the PDGFRβ signaling experiment, cells were serum-starved overnight following transfection and then stimulated with recombinant human PDGF-BB (rhPDGF-BB) for 30 minutes at the indicated concentrations, and the cells were stored in -80 °C for subsequent experiments.

### Western blot and immunoprecipitation analysis

HEK-293T cells were washed once with PBS and lysed on ice for 5 minutes in lysis buffer containing 25 mM HEPES (pH 7.4), 150 mM NaCl, 5 mM EDTA, 10 µg/ml aprotinin, 5 µg/ml leupeptin, 10% glycerol, and 1% Triton X-100. Cell lysates were clarified by centrifugation at 13,500 × g for 20 minutes at 4°C. The supernatants were then incubated with 1 µg of the indicated antibody and 20 µl of 50% (v/v) Protein A-agarose (Cytiva, Marlborough, MA, USA) for 2 hours with gentle rocking. After three washes with lysis buffer, the immunoprecipitated complexes were solubilized by boiling in Laemmli sample buffer, fractionated by SDS-PAGE, and transferred onto PVDF membranes. The membranes were blocked with PBS (pH 7.4) containing 5% BSA and 0.05% Tween 20, followed by incubation with the indicated primary antibodies. After three additional washes, the membranes were incubated with HRP-conjugated secondary antibodies for 1 hour. Following three final washes, protein bands were visualized using the VisGlow chemiluminescent HRP substrate system (Visual Protein, Taipei, Taiwan).

## Results

### Dormant expression of Scube2 in epicardial tissue and reactivation upon cardiac cryoinjury

To investigate the expression of *scube2* in adult hearts, we performed *in situ* hybridization on adult heart sections with and without cryoinjury (Figure 1). We found *scube2* was expressed mainly in the epicardial tissue of uninjured hearts and reactivated at both 3 and 7 days post-cryoinjury (dpci; Figure 1A-C and Figure S1A and B). To confirm the expression of Scube2 in the epicardium, we performed immunostaining for Scube2 under various transgenic reporter backgrounds. Immunostaining results further confirmed the reactivation of Scube2 in epicardial cells, which colocalized with epicardial *Tg(tcf21:NLS-EGFP)* and *Tg(wt1b:EGFP)*, but not endothelial *Tg(fli1:EGFP)* reporter expression (Figure 1D-H). This tissue-specific expression and reactivation of *scube2* in the epicardial tissue of zebrafish hearts correspond nicely to published bulk and single-cell RNAseq datasets (35, 36). Furthermore, *scube2* reactivation is absent in another teleost model, the medaka, which cannot regenerate their hearts post-cryoinjury (Figure S1B)(28, 35). Altogether, these findings suggest a correlation of Scube2 expression with cardiac development and regeneration after cardiac injury.

**Figure 1.**
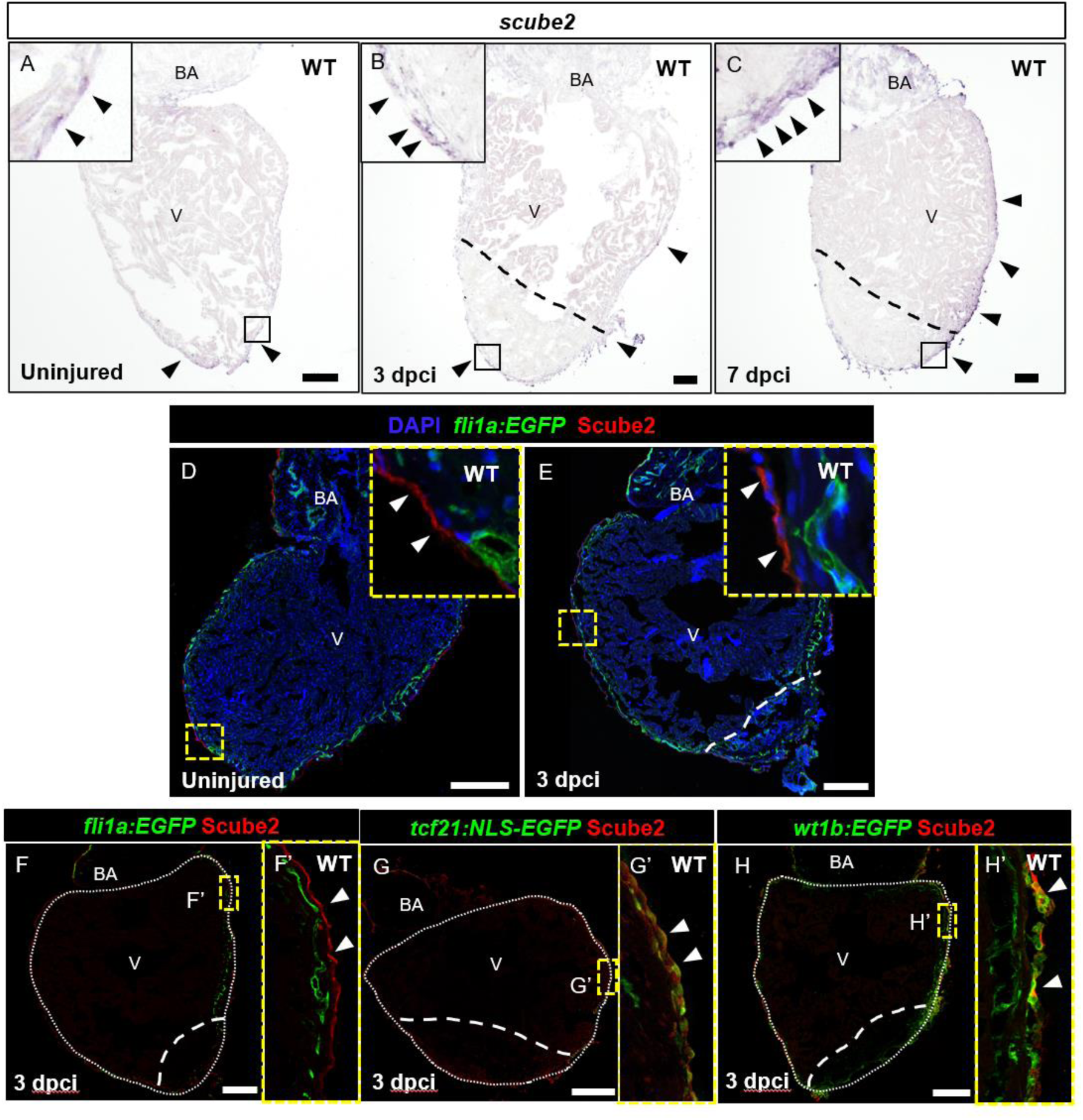
Scube2 expression in the epicardium of homeostatic and injured hearts. (A–C) In situ hybridization (ISH) of *scube2* in uninjured, 3 days post cardiac injury (3 dpci), and 7 dpci zebrafish hearts. Expression of *scube2* (arrowheads) was weakly detected in the outer layer of the ventricle under homeostatic conditions and was reactivated following cardiac injury. Scale bar = 100 μm. (D–F) Immunohistochemistry (IHC) for Scube2 in uninjured and 3 dpci hearts in the *Tg(fli1a:EGFP)*. (G) IHC for Scube2 in the *Tg(tcf21:NLS-EGFP)* background. (H) IHC for Scube2 in the *Tg(wt1b:EGFP)* background. (G′, H′) Insets show epicardial-derived cells expressing Scube2 (arrowheads). Scale bar = 200 μm.

### Scube2 regulates coronary vessel formation during cardiac growth

To assess the role of Scube2 in coronary development, we analyzed the coronary vasculature of *scube2* mutants (*scube2^Δ1/Δ1^*) and compared it to that of wild-type (WT) fish in the *Tg(fli1a:EGFP)* background at different stages of coronary network formation (Figure 2). Confocal microscopy revealed significantly reduced coronary network coverage in the ventricles of in *scube2^Δ1/Δ1^* compared to WT at 17 mm length (body length from the snout to the base of the tail fin (Figure 2A and B, quantified in E). Coronary vasculature was observed in both *scube2^Δ1/Δ1^*and WT ventricles at 28 mm, but coronary network coverage remained reduced when compared to WT ventricles (roughly 7-8 mpf, Figure 2C and D, quantified in F). At this stage (>6 months old), we stained the ventricle sections with Toluidine blue and observed necrotic myocardium and reduced coronary vessels in *scube2^Δ1/Δ1^* hearts compared to WT hearts (Figure 2G-H). Electron microscopy further revealed disorganized myofibrils (Figure 2I-J and L-M) and damaged mitochondria (Figure 2K and N) in *scube2^Δ1/Δ1^* hearts, presumably a secondary effect of deformed/shrunken coronary vessels (Figure 2K and N). Consistent with the observed role of Scube2 in coronary development in zebrafish, the expression level of *Scube2* peaks at the coronary establishment and expansion stages in both epicardium and endothelium (Figure S2). In mouse hearts, the *Scube2* expression is also high during embryonic stages and declines after birth when the coronary vasculature matures (Figure S3). Interestingly, P8 is also the stage when neonatal mice lose regenerative capacity (37). These results support the role of Scube2 in coronary vessel formation during cardiac growth.

**Figure 2.**
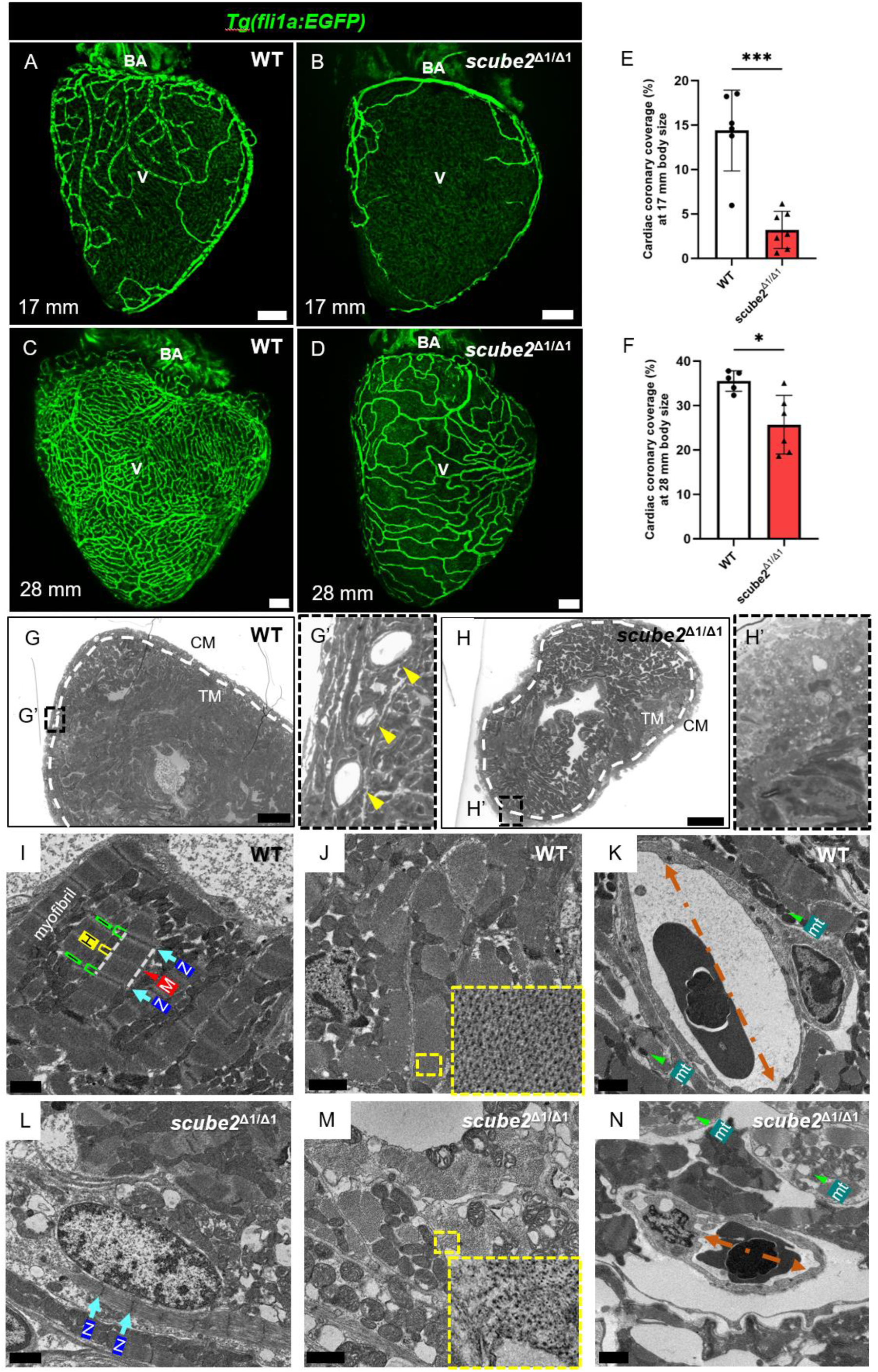
Defective coronary vasculature and myocardium in *scube2* mutants. (A–D) Confocal images of the coronary vasculature in wild-type (WT) and *scube2* mutant zebrafish hearts at the developing stage (17 mm snout-fin base length) and maturation stage (28 mm snout-fin base length). BA: bulbus arteriosus; V: ventricle. Ventral view. Scale bar = 100 µm. (E, F) Quantification of coronary vessel coverage at the developing and maturation stages using ImageJ. Data were analyzed using Student’s t-test (developing stage: n ≥ 7, ***P = 0.0001; maturation stage: n ≥ 6, *P < 0.05). (G, H) Ventricle cross-sections and Toluidine Blue staining from adult WT and *scube2* zebrafish. The compact myocardium (outlined by dotted lines) and coronary vessels (yellow arrowhead, G′) are indicated. CM: cortical myocardium; TM: trabecular myocardium. Scale bar = 100 µm. (I–N) Transmission electron microscopy (TEM) images of cardiac ultrastructure in WT and *scube2* mutant adult hearts. Scale bar = 1 µm. (I, L) WT myofibrils show clearly defined sarcomeres, marked by distinct Z-lines and M-lines, whereas *scube2* mutants exhibit necrotic myocardium with disorganized myofibrils. Scale bar = 1 µm. (J, M) The myofibrillar arrangement in cross-sections is irregular and disorganized in *scube2* mutants compared to WT. Scale bar = 1 µm. (K, N) Coronary vessels in *scube2* mutants are more constricted and deformed compared to WT. Degradation of surrounding mitochondria (mt) is also evident in mutants, in contrast to the intact mitochondria in WT hearts. Scale bar = 1 µm.

### Scube2 deficiency impairs revascularization and cardiac regeneration, leading to unresolved scar post cardiac injury

To investigate whether Scube2 is involved in cardiac regeneration, we performed cardiac cryoinjury on *scube2^Δ1/Δ1^*, *scube2^Δ1/+^*, and WT fish, and examined the regeneration processes at various time points after cryoinjury (20, 21). The injury border zone has been shown to be the regenerative niche where the mature cardiomyocytes re-enter the cell cycle and proliferate for regeneration (38, 39). Consistent with previous observations (9), WT hearts formed a new vascular network that supports tissue regeneration along with the resolution of fibrotic tissue by 30 dpci (Figure 3B, E, H). On the contrary, both *scube2^Δ1/+^*(Figure 3C, F, I) and *scube2^Δ1/Δ1^* (Figure 3D, G, J) hearts exhibited reduced vascularized areas at 7 (Figure 3K) and 30 dpci (Figure 3L).

**Figure 3.**
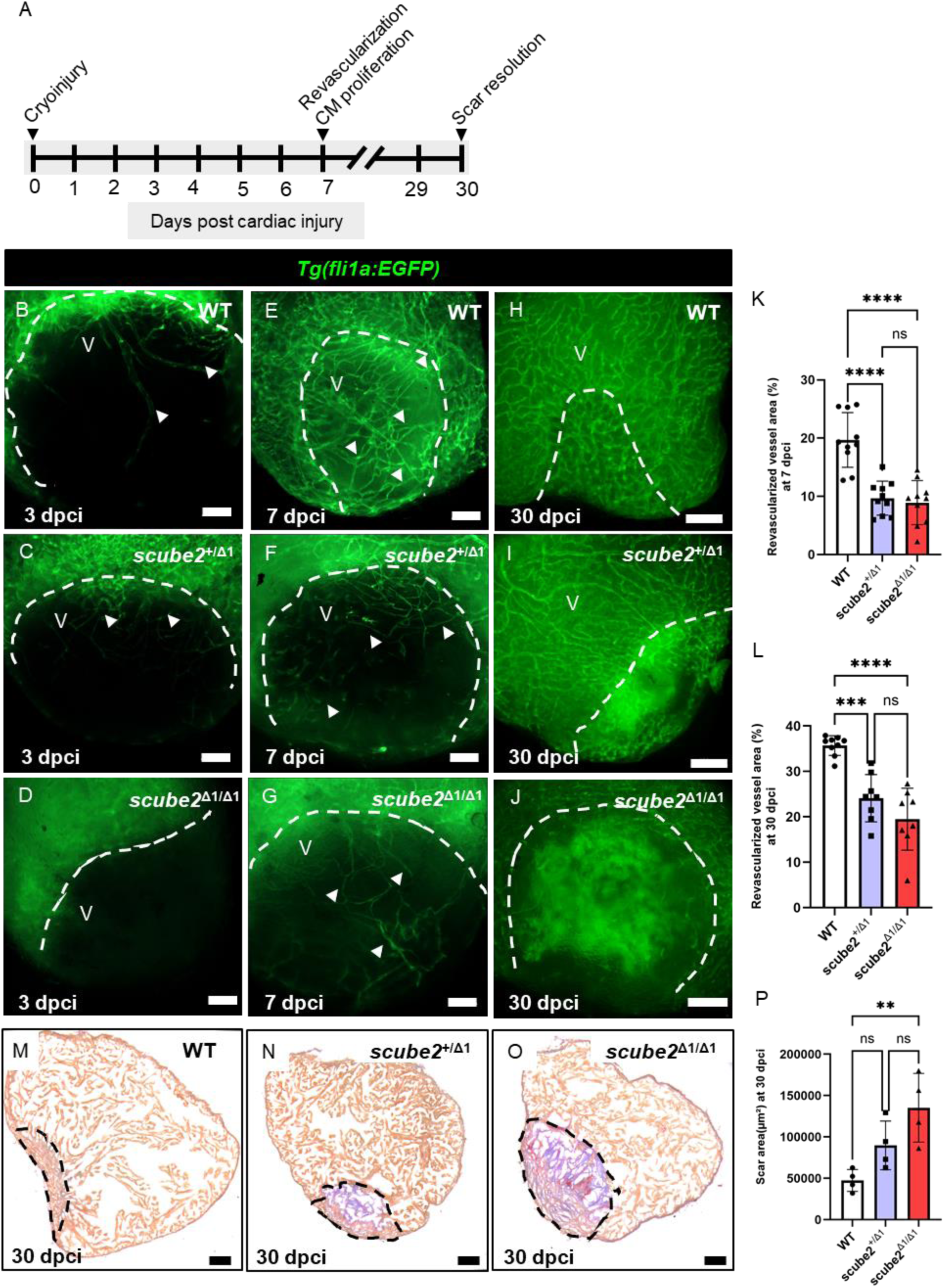
Loss of *scube2* impairs cardiac revascularization and scar resolution during cardiac repair. (A–J) Cryoinjured hearts from wild-type, heterozygous, and homozygous *scube2* mutants in the Tg(*fli1a:EGFP*) background were collected at 3, 7, and 30 days post cardiac injury (dpci). Scale bars = 100 μm. (K) Quantification of revascularization at 7 dpci using ImageJ. Data were analyzed by one-way ANOVA (*n* = 10; \*\*\**P* < 0.0001). Error bars represent ± standard deviation. (L) Quantification of revascularization at 30 dpci using ImageJ. Data were analyzed by one-way ANOVA (*n* ≥ 8; \*\*\**P* < 0.0001; \*\**P* < 0.001). Error bars represent ± standard deviation. (M–O) Cryosections of WT and *scube2* mutant hearts at 30 dpci stained with Acid Fuchsin Orange G (AFOG) to label collagenous scar tissue. Scale bars = 100 μm. (P) Quantification of collagenous scar area at 30 dpci using NIS-Elements AR 5.21.03 software, analyzed by one-way ANOVA (*n* = 4; \**P* < 0.01). Error bars represent ± standard deviation.

To examine the outcome of cardiac repair, we performed AFOG staining on cryosections of *scube2* mutant and WT hearts at 30 dpci, in which collagen tissue will be stained in blue, fibrin in red, and the myocardium in orange (Figure 3M-O, quantified in P). While the fibrotic tissues were much reduced and replaced by regenerated myocardium in WT hearts, *scube2^Δ1/Δ1^* hearts showed significantly larger collagen deposits, corresponding to the observations under a fluorescent microscope with *Tg(fli1a:EGFP)* reporter background. These results indicate that Scube2 is essential for cardiac regeneration.

### Scube2 modulates VEGF and PDGF signaling during cardiac regeneration

To investigate the molecular mechanisms underlying Scube2 function in cardiac regeneration, we collected *scube2^Δ1/Δ1^* and WT hearts before injury and at 3 and 7 dpci for RNAseq analyses (Figure 4A). PCA analysis revealed a similar trend of dynamic changes in response to injury in both *scube2^Δ1/Δ1^* and WT hearts, as indicated by PC1. However, recovery of *scube2^Δ1/Δ1^* hearts from 3 to 7 dpci was less compared to WT hearts. In PC2, the data points for *scube2^Δ1/Δ1^*and WT hearts were completely separated, corresponding to the intrinsic defects observed in untreated hearts (Figure 4B).

**Figure 4.**
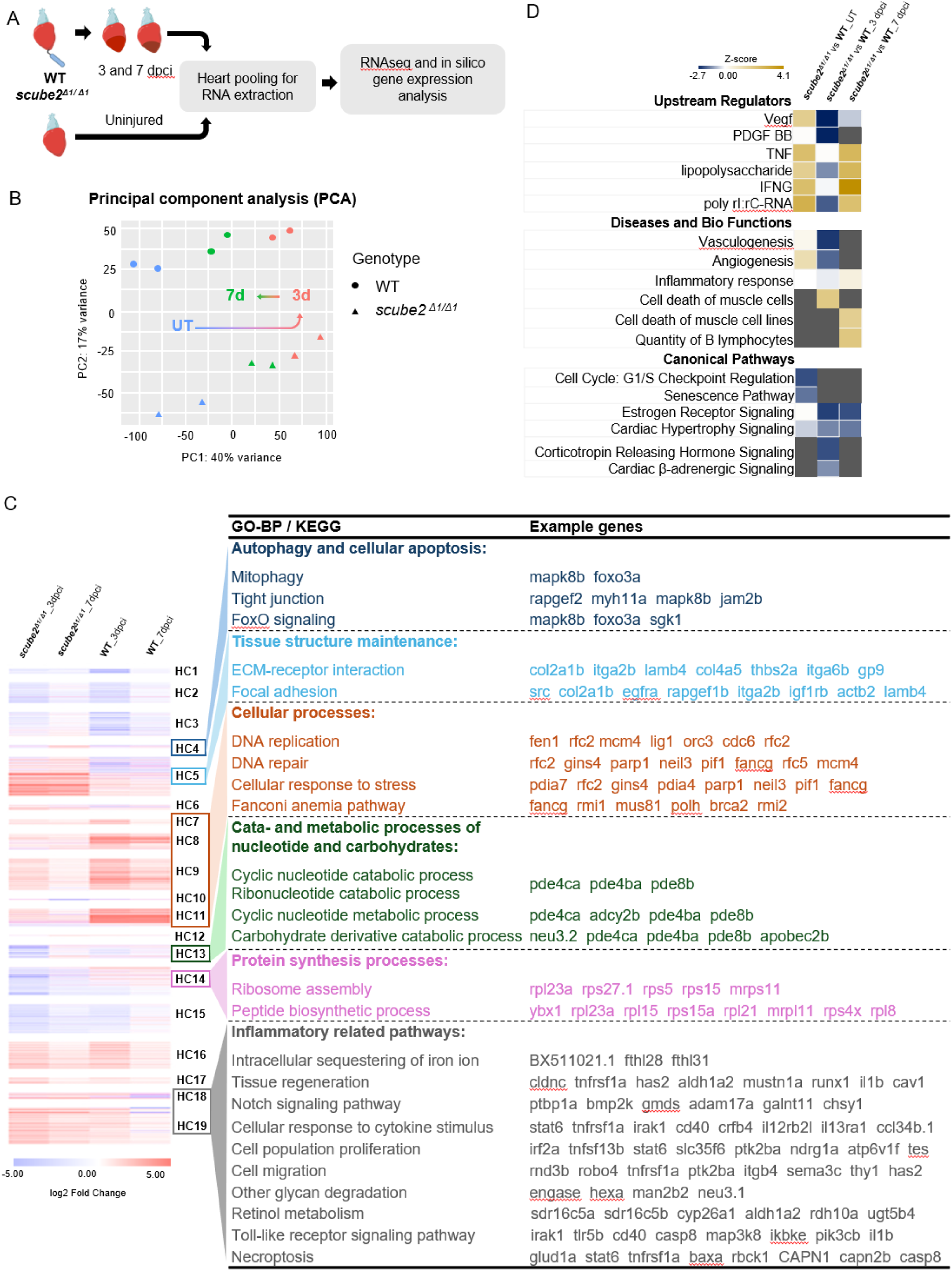
Transcriptional profiling of injured hearts in *scube2* mutant versus wild-type zebrafish. (A) Hearts from wild-type and scube2 mutant zebrafish were collected before injury (0 dpci) and at 3 and 7 days post cardiac injury (dpci), pooled for RNA extraction, and subjected to next-generation sequencing. (B) Principal component analysis (PCA) of the transcriptomic data revealed distinct differences between wild-type and scube2 mutant hearts at the transcriptomic level. (C) Hierarchical clustering of differentially expressed genes showed that scube2 mutant hearts had increased expression of genes related to apoptosis and inflammation, suggesting persistent immune activation and impaired resolution, potentially contributing to larger scar formation. In contrast, genes associated with DNA replication were more highly expressed in wild-type hearts, indicating greater proliferative activity—possibly in cardiomyocytes or endothelial cells—consistent with enhanced revascularization and myocardial regeneration. Genes involved in protein biosynthesis and metabolic processes were broadly downregulated, aligning with impaired repair capacity observed in scube2 mutants. (D) Ingenuity Pathway Analysis (IPA) was used to identify upstream regulators, associated diseases, biofunctions, and canonical pathways. The analysis was filtered to include terms with – log(p-value) > 1.5.

We next performed hierarchical clustering of differentially expressed genes (DEGs) across genotypes and time points and ran Gene Ontology (GO) analyses to explore their biological functions (Figure 4C). DEGs associated with “autophagy”, “cellular apoptosis”, and “tissue structure maintenance” were preferentially activated in mutants compared to the WT, suggesting that excessive cell death and stress occurred in mutant conditions and required extra help in restoring tissue integrity (HC4 and 5). On the other hand, WT preferentially activated DEGs associated with various aspects of tissue repair, including DNA replication and repair, metabolic and catabolic processes, and protein synthesis, which correlates with generating new building blocks for cell replenishment (HC7-14).

Lastly, we noticed DEGs related to inflammatory pathways remained active at 7 dpci in mutant hearts, suggesting a prolonged inflammation potentially associated with excessive cell death and impaired tissue repair (HC18-19).

Next, we performed Ingenuity Pathway Analysis to identify potential pathways and biological functions directly associated with *scube2* deficiency during cardiac repair (Figure 4D). Among upstream regulators, VEGF and PDGF were differentially enriched in WT compared to mutant hearts, while inflammatory cytokines, such as TNF and IFNG), show higher levels at homeostasis and 7 dpci. These observations nicely correlate with the biological functions of vasculogenesis/angiogenesis in WT hearts and inflammatory cell death in mutant hearts. Furthermore, canonical pathways previously described in heart regeneration, including cell cycle regulation (40), estrogen (41), and endocrine hormones (42), were preferentially enriched in WT compared to *scube2* mutants. These results align well with our histological examination and functional investigations, as well as previous studies, supporting the role of Scube2 in modulating vascular formation and heart regeneration through VEGF and PDGF signaling.

### SCUBE2 enhances PDGF signaling via interaction with PDGF-B and its Receptor

VEGF and PDGF function as potent mitogens for both endothelial cells and cardiomyocytes during development and regeneration with potential crosstalk (9, 43–48). During cardiac repair, revascularization of the infarct and restoration of myocardial tissue and contractile function also rely on the replenishment of vascular endothelial cells and cardiomyocytes (9, 18, 40). While Scube2 acts as a co-receptor in accelerating VEGF signaling, its potential function in PDGF signaling is uncharacterized.

To determine whether SCUBE2 enhances PDGF signaling, we first examined its potential interaction with PDGF-B. Co-immunoprecipitation (co-IP) assays in HEK-293T cells co-transfected with FLAG-tagged SCUBE2 and His-tagged PDGF-B revealed that SCUBE2 physically associates with PDGF-B, as evidenced by the co-precipitation of FLAG.SCUBE2 using an anti-His antibody (Figure 5A, upper panel). Protein expression was confirmed by Western blotting (WB) using anti-FLAG and anti-His antibodies (Figure 5A, middle and lower panels).

**Figure 5.**
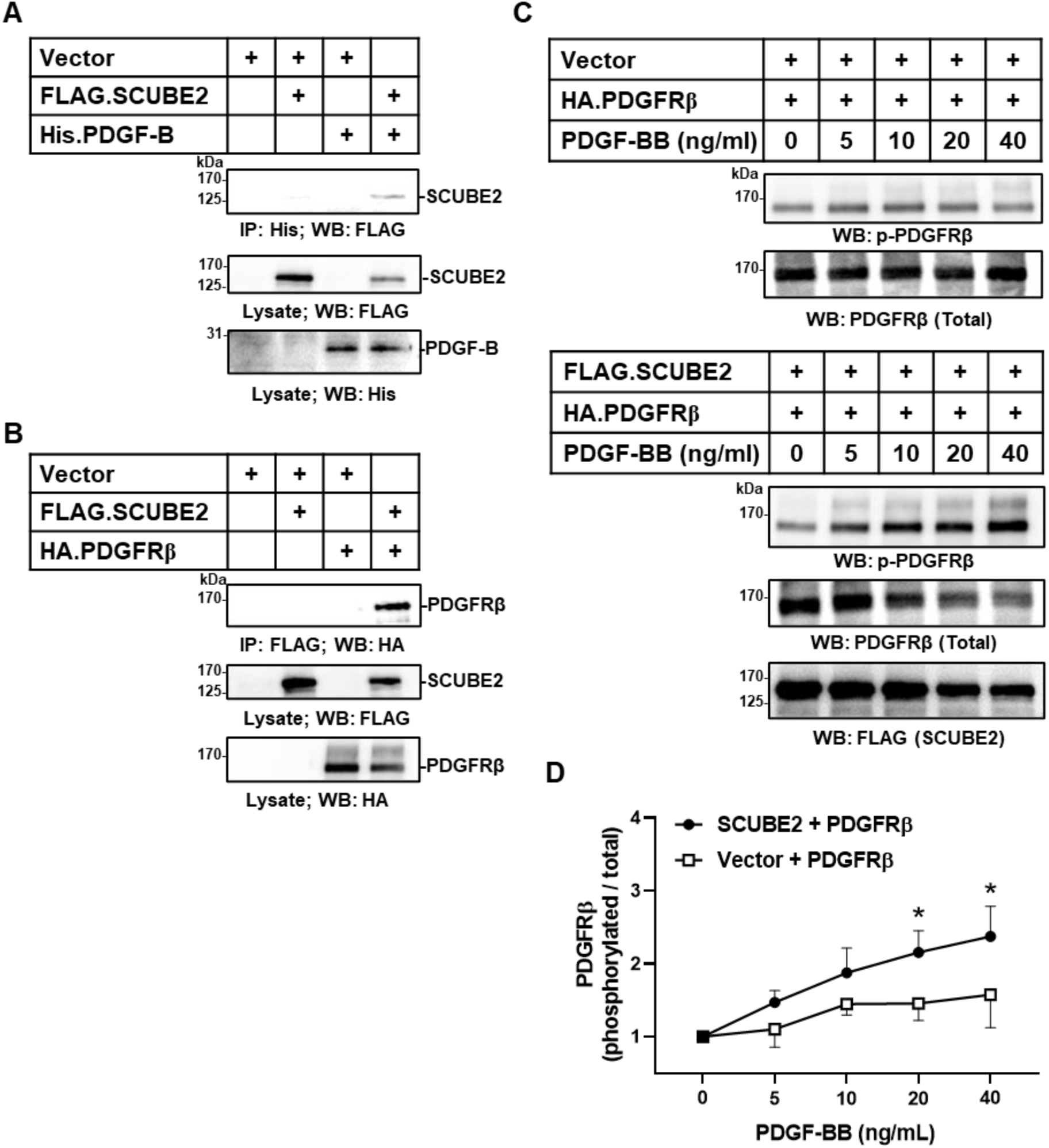
SCUBE2 enhances PDGF signaling via interaction with PDGF-B and its receptor. (A) SCUBE2 interacts with the PDGF-B ligand. HEK-293T cells were transfected with FLAG-tagged SCUBE2 (FLAG.SCUBE2) and His-tagged PDGF-B (His.PDGF-B) individually or in combination for two days. Cell lysates were subjected to immunoprecipitation (IP) using an anti-His antibody, and the co-immunoprecipitated SCUBE2 protein was detected by Western blot (WB) analysis using an anti-FLAG antibody (upper panel). The expression of FLAG.SCUBE2 and His.PDGF-B was confirmed by anti-FLAG and anti-His WB analysis, respectively (middle and lower panels). (B) SCUBE2 is capable of binding PDGFRβ. HEK-293T cells were transfected with FLAG.SCUBE2 and HA-tagged PDGFRβ (HA.PDGFRβ) either individually or together for two days. Lysates were immunoprecipitated using an anti-FLAG antibody, and the associated PDGFRβ protein was detected by WB using an anti-HA antibody (upper panel). The expression of FLAG.SCUBE2 and HA.PDGFRβ was validated by anti-FLAG and anti-HA WB analysis, respectively (middle and lower panels). (C) SCUBE2 promotes PDGF-BB-induced PDGFRβ signaling. HEK-293T cells were co-transfected with vector control and FLAG.SCUBE2, or with HA.PDGFRβ and FLAG.SCUBE2, for two days. PDGFRβ signaling was then stimulated by the addition of recombinant PDGF-BB protein for 30 minutes at the indicated concentrations. The levels of phosphorylated PDGFRβ (p-PDGFRβ) as indication of ligand-induced signaling, total protein of PDGFRβ, and SCUBE2 were analyzed by WB using anti-p-PDGFRβ, anti-PDGFRβ, and anti-FLAG antibodies, respectively. Protein levels of p-PDGFRβ and total PDGFRβ were quantified and plotted as a line graph (D). Statistical comparison was performed between the SCUBE2 + PDGFRβ and vector + PDGFRβ groups. Data are presented as mean ± S.D. from four independent experiments. *, *P* < 0.05.

Next, we tested whether SCUBE2 also binds the PDGF receptor β (PDGFRβ). Co-IP analysis in HEK-293T cells co-expressing FLAG.SCUBE2 and HA-tagged PDGFRβ demonstrated that SCUBE2 interacts with PDGFRβ. Specifically, HA.PDGFRβ was detected in anti-FLAG immunoprecipitates only when both proteins were co-expressed (Figure 5B, upper panel), confirming their association. The expression of FLAG.SCUBE2 and HA.PDGFRβ was validated by WB (Figure 5B, middle and lower panels).

To assess the functional consequence of these interactions, we examined PDGF-BB-induced PDGFRβ activation in cells expressing SCUBE2. Upon stimulation with recombinant PDGF-BB for 30 minutes, cells co-expressing FLAG.SCUBE2 and HA.PDGFRβ exhibited significantly increased phosphorylation of PDGFRβ (p-PDGFRβ) compared to controls, indicating enhanced receptor activation (Figure 5C). Quantification of p-PDGFRβ relative to total PDGFRβ levels showed a statistically significant increase in ligand-induced signaling in the SCUBE2 + PDGFRβ group compared to the vector + PDGFRβ group (Figure 5D). These results suggest that SCUBE2 potentiates PDGF signaling by interacting with both PDGF-B and PDGFRβ.

### Scube2 promotes endothelial and cardiomyocyte proliferation during cardiac regeneration

Enlightened by the potential role of Scube2 in accelerating VEGF and PDGF signaling during cardiac regeneration, we further examined whether Scube2 regulates the proliferation and replenishment of endothelial cells and CMs during cardiac regeneration. To this end, we performed cryoinjury, collected the hearts at 7 dpci, and performed PCNA staining under the *Tg(fli1a:EGFP)* and *Tg(myl7:nuc-RFP)* reporter backgrounds, respectively (Figure 6). In agreement with our hypothesis, both *scube2^Δ1/Δ1^* and *scube2^Δ1/+^* hearts showed significantly lower amounts of CMs proliferation than WT hearts (Figure 5A-C, quantified in D). Similarly, *scube2^Δ1/Δ1^* hearts showed significantly lower amounts of endothelial cell proliferation than WT hearts (Figure 5E-G, quantified in H). Aligning with results from *in silico* and biochemical analyses, these data support the role of Scube2 in modulating endothelial cell and CM proliferation during revascularization and heart regeneration.

**Figure 6.**
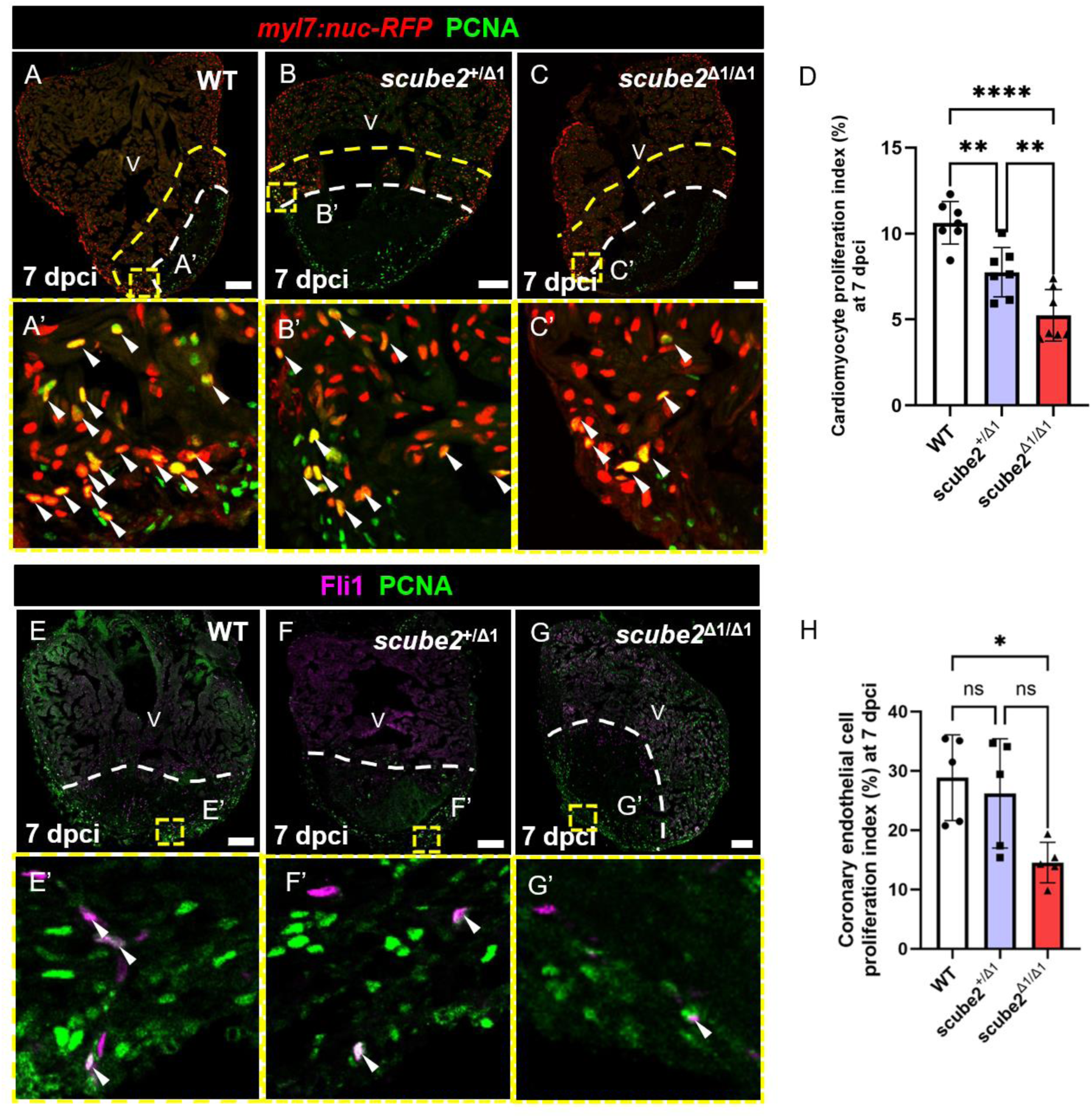
Loss of *scube2* impairs the proliferation of cardiomyocytes and endothelial cells during cardiac repair. (A–C) Cryosections of *scube2* mutant and wild-type (WT) hearts at 7 days post cryoinjury (dpci) were immunostained with proliferating cell nuclear antigen (PCNA) and anti-mCherry antibody to detect proliferating cardiomyocytes in the *myl7:nuc-RFP* background. Insets (A′–C′) highlight proliferating cardiomyocytes (arrowheads). Scale bars = 100 μm. (D) Quantification of proliferating border zone cardiomyocytes at 7 dpci (*n* = 7; \**P* < 0.01, \*\*\**P* < 0.0001). Error bars represent ± standard deviation. (E–G) Cryosections of *scube2* mutant and WT hearts at 7 dpci were immunostained with PCNA and anti-Fli1 antibody to detect proliferating coronary endothelial cells (arrowheads). Insets (E′–G′) highlight proliferating endothelial cells. Scale bars = 100 μm. (H) Quantification of proliferating coronary endothelial cells at 7 dpci (*n* = 5; *P* < 0.05). Error bars represent ± standard deviation.

## Discussion

Our study identified a novel role of Scube2 in the formation of the coronary vasculature during zebrafish cardiac development and regeneration. *scube2* mutants exhibit impaired coronary vasculature, highlighting its role in physiological angiogenesis of the heart. Furthermore, defects in the myocardium were observed in *scube2* mutants during homeostasis and after cardiac injury, suggesting a contribution of coronary vessels to CM growth and regeneration, regulated by Scube2.

In addition, we found that *scube2* is expressed in the epicardium and upregulated after cryoinjury. The epicardium has been considered a signaling hub for heart regeneration, serving as a source of crucial cells, including vascular smooth muscle cells, pericytes, and fibroblasts, and secreting factors that are essential for the proliferation and survival of cardiomyocytes (49). Indeed, transcriptomic analyses revealed disrupted VEGF and PDGF signaling pathways, as well as persistent inflammation in *scube2*-deficient hearts, correlating with impaired vascular and regenerative responses. Functional experiments revealed impaired proliferation of both CMs and endothelial cells, which is closely associated with defects in revascularization and the replacement of scar tissue with new CMs. In addition, genes associated with cell cycle, sarcomere assembly, and hormonal signaling were dysregulated, indicating that Scube2 may influence cardiac regeneration beyond angiogenesis, possibly affecting cardiomyocyte architecture and signaling milieu. Lastly, although we did not observe related changes in the RNA-seq profiling results, the Hh signaling pathway is also regulated by Scube2 during embryonic development and is associated with coronary formation (50) and CM proliferation (51, 52) during cardiac repair.

These findings position Scube2 as a key regulator of coronary development and heart regeneration. The evolutionary conservation of its signaling enhancer role, along with its preferential expression in regenerative zebrafish compared to non-regenerative medaka and adult mice, hints at therapeutic potential in accelerating cardiac repair. Future work should explore the gain-of-function and delivery strategies for Scube2 and further dissect the interaction between Scube2-mediated angiogenesis and other signaling pathways during heart regeneration.

## Conclusion

In summary, this study identifies Scube2 as a novel regulator of coronary vessel development and heart regeneration in zebrafish. Scube2 functions through VEGF signaling enhancement, contributing to both physiological angiogenesis and injury-induced revascularization. Loss of Scube2 disrupts vascular growth, impairs CM replenishment, and leads to persistent inflammation and fibrotic scarring. These findings suggest the potential of targeting Scube2 to enhance coronary vascularization and myocardial regeneration in ischemic heart disease.

## Acknowledgments

We thank the core facilities at the Institute of Biomedical Sciences, including Light Microscopy, Pathology and DNA Sequencing Core; Dr. Yao-Ming Chang and the Computational Medicine Core; Dr. Meiyeh Jade Lu and the High Throughput Genomics Core at Biodiversity Research Center for the NGS work; Mr. Yao-Kwan Huang and the Electron Microscopy Core facility at Institute of Cellular and Organismic Biology of Academia Sinica. We are also grateful for the technical services provided by the Transgenic Mouse Model Core Facility of the National Core Facility Program for Biotechnology, National Science and Technology Council (NSTC) of Taiwan, and the Gene Knockout Mouse Core Laboratory of the National Taiwan University Center of Genomic Medicine for help and advice in producing the knockout mice.

---

**Figure S1.**
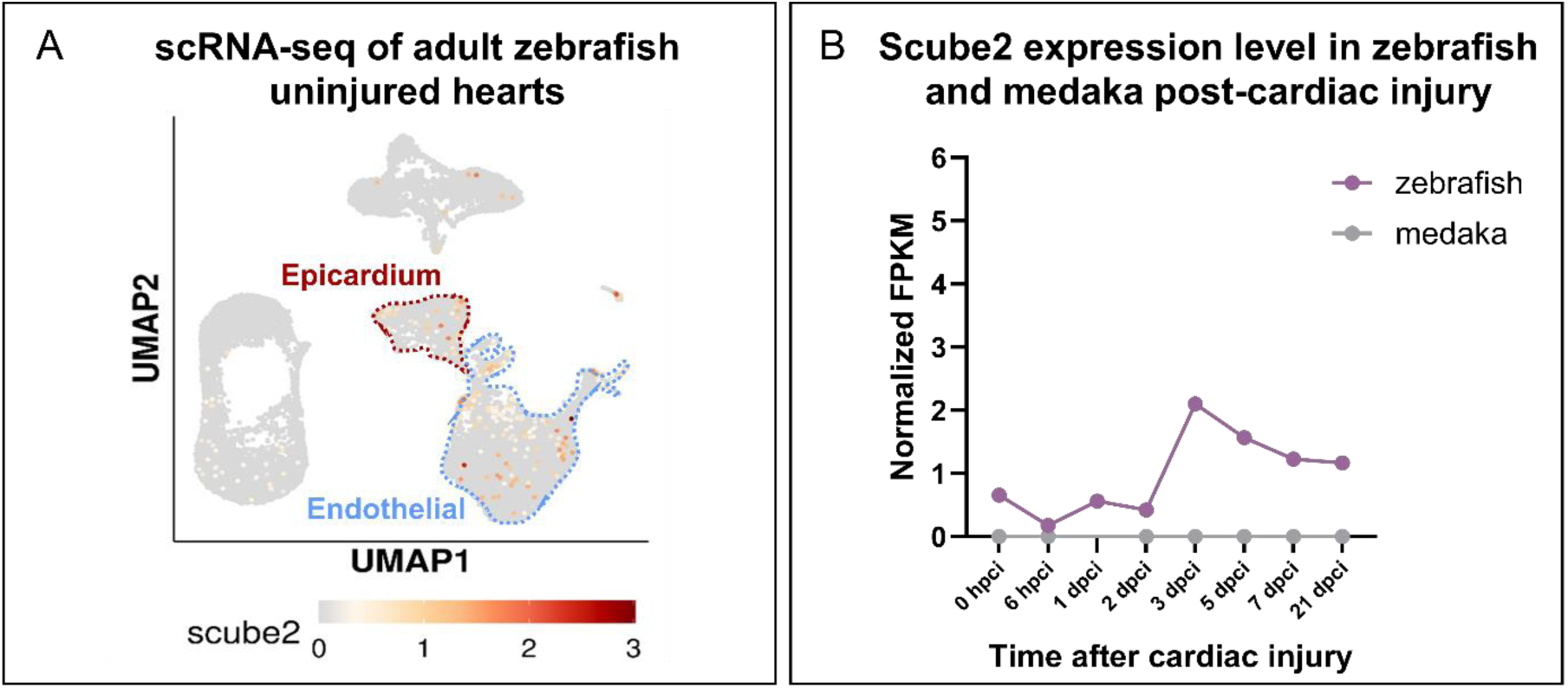
In silico analyses of *scube2* expression in adult zebrafish and medaka hearts before and after injury. (A) Expression of *scube2* in the epicardium and endothelial cells of adult zebrafish uninjured hearts was extracted from a published single-cell RNA-sequencing dataset (Carey CM et al., 2024). (B) Expression data of *scube2* genes in adult (regenerative) zebrafish and (non-regenerative) medaka hearts before and following injury were extracted from the bulk RNAseq (Wei et al., 2023, and unpublished data, Chowdhury K et al.).

**Figure S2.**
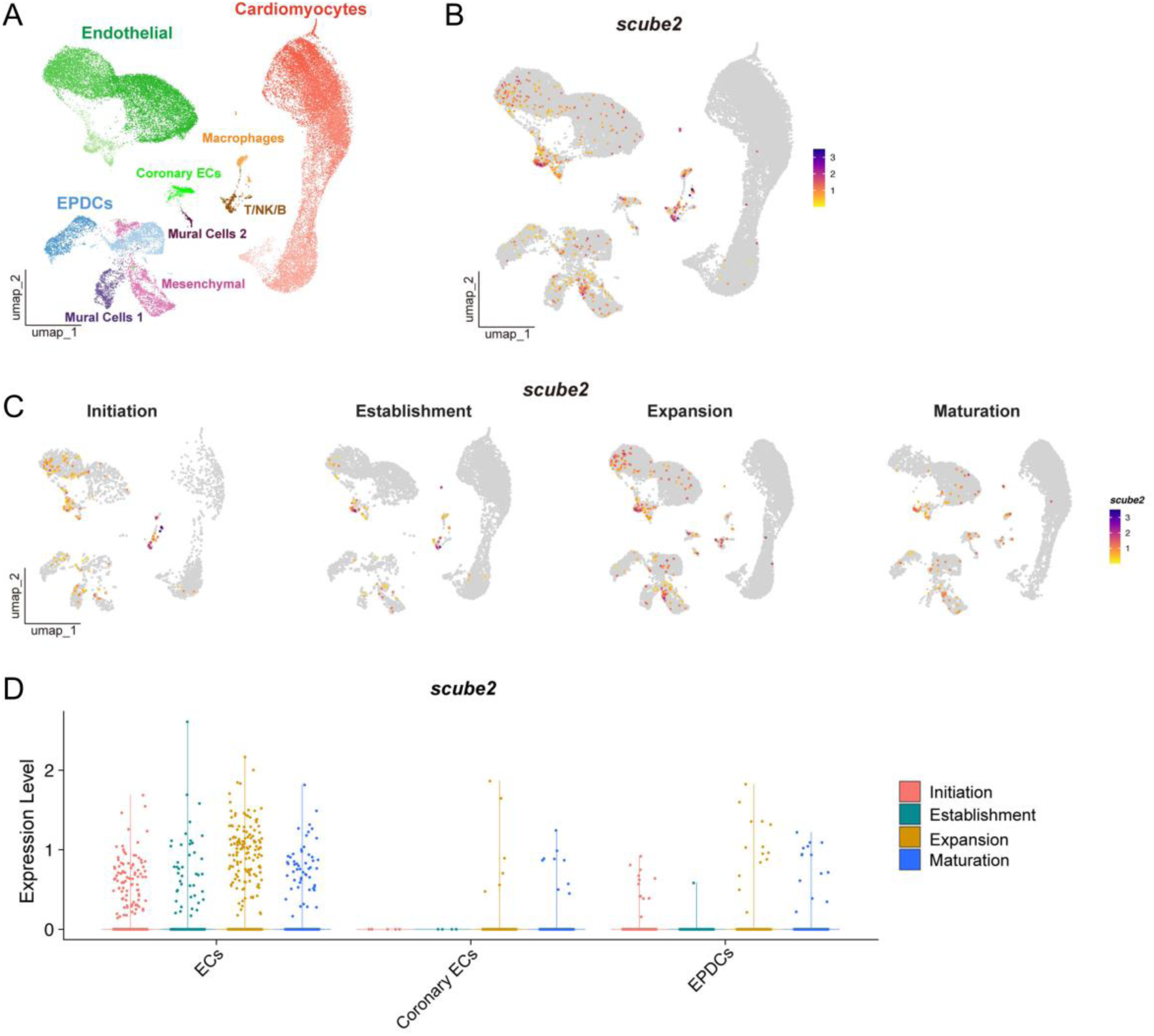
Expression of *scube2* in the zebrafish heart during juvenile and adult stages, corresponding with coronary vascular development. (A) UMAP embedding the main cardiac cell types of zebrafish developmental hearts. Data was extracted from single-cell RNAseq experiments in Rouf et al. (under revision). (B) Expression of *scube2* was most abundant in endocardial cells (ECs), coronary endothelial cells, and epicardial-derived cells (EPDCs). (C and D) Corresponding to coronary vessel development, *Scube2* expression peaks between the establishment and expansion phases, before decreasing again in the maturation phase.

**Figure S3.**
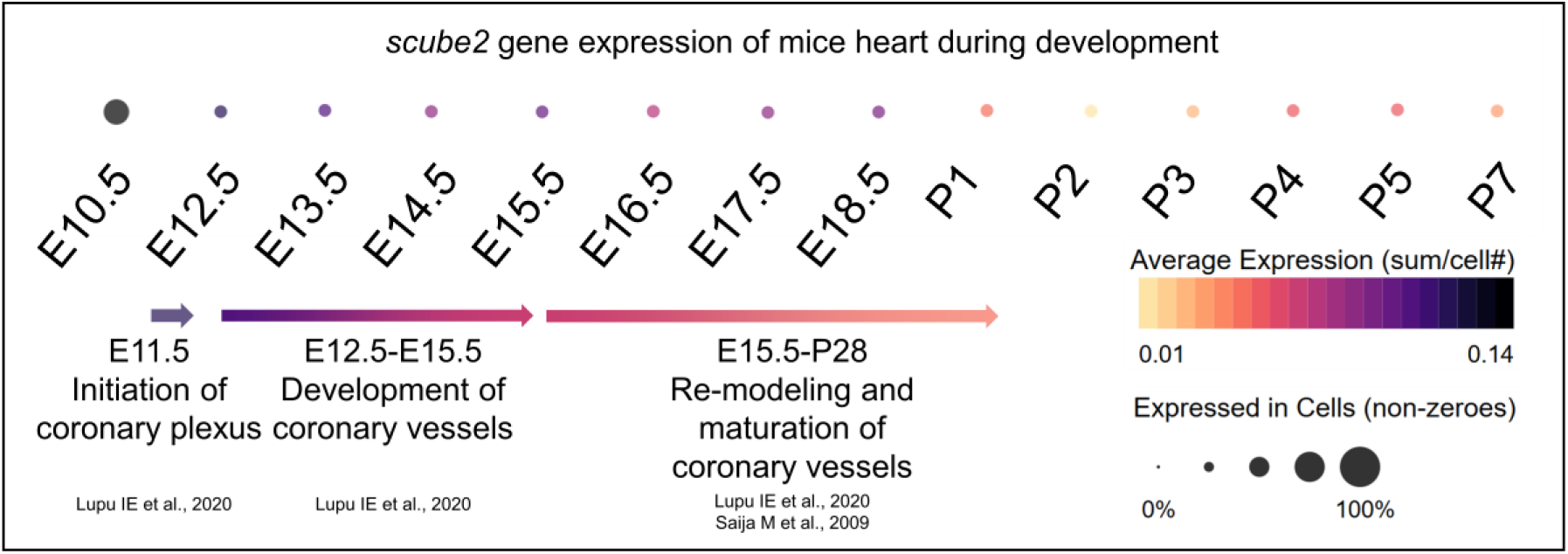
Expression of *Scube2* in the mouse heart during embryonic and postnatal stages, corresponding with coronary vascular development. *Scube2* expression is elevated during embryonic stages and declines after the formation of the coronary vasculature in postnatal mouse hearts. Timeline of coronary development was referenced from <u>Saija M et al., 2009</u> and <u>Lupu IE et al., 2020</u>; Expression data of Scube2 was extracted from the single-cell RNA-sequencing experiment from <u>Feng W et al., 2022</u>. Average expression of *Scube2* across all cell types was analyzed to assess its association with coronary development, since Scube2 encodes a secreted protein.

